# Mouse hepatitis virus nsp14 exoribonuclease activity is required for resistance to innate immunity

**DOI:** 10.1101/182196

**Authors:** James Brett Case, Yize Li, Ruth Elliott, Xiaotao Lu, Kevin W. Graepel, Nicole R. Sexton, Everett Clinton Smith, Susan R. Weiss, Mark R. Denison

## Abstract

Coronaviruses (CoV) are positive-sense RNA viruses that infect numerous mammalian and avian species and are capable of causing severe and lethal disease in humans. CoVs encode several innate immune antagonists that interact with the host innate immune response to facilitate efficient viral replication. CoV non-structural protein 14 (nsp14) encodes 3'-to-5' exoribonuclease activity (ExoN), which performs a proofreading function and is required for high-fidelity replication. Outside of the order *Nidovirales*, arenaviruses are the only RNA viruses that encode an ExoN, which functions to degrade dsRNA replication intermediates. In this study, we tested the hypothesis that CoV ExoN may also function to antagonize the innate immune response. We demonstrate that viruses lacking ExoN activity [ExoN(-)] are sensitive to cellular pretreatment with interferon beta (IFN-β) in a dose-dependent manner. In addition, ExoN(-) virus replication was attenuated in wild-type bone marrow-derived macrophages (BMMs) and partially restored in interferon alpha/beta receptor deficient (IFNAR-/-) BMMs. ExoN(-) virus replication did not result in IFN-β gene expression, and in the presence of an IFN-β-mediated antiviral state, ExoN(-) viral RNA levels were not substantially reduced relative to untreated. However, ExoN(-) virus generated from IFN-β pretreated cells had reduced specific infectivity and decreased relative fitness, suggesting that ExoN(-) virus generated during an antiviral state is less viable to establish a subsequent infection. Overall, our data suggest MHV ExoN activity is required for resistance to the innate immune response and antiviral mechanisms affecting the viral RNA sequence and/or an RNA modification act on viruses lacking ExoN activity.

## IMPORTANCE

CoVs encode multiple antagonists that prevent or disrupt an efficient innate response. Additionally, no specific antiviral therapies or vaccines currently exist for human CoV infections. Therefore, the study of CoV innate immune antagonists is essential for understanding how CoVs overcome host defenses and to maximize potential therapeutic interventions. Here, we sought to determine the contributions of nsp14 ExoN activity in the induction of and resistance to the innate immune response. We show that viruses lacking nsp14-ExoN activity are more sensitive to restriction by exogenous IFN-β and that viruses produced in the presence of an antiviral state are less capable of establishing a subsequent viral infection. Our results support the hypothesis that MHV ExoN activity is required for resistance to the innate immune response.

## INTRODUCTION

The innate immune response within a mammalian cell is the first line of defense against an invading pathogen. However, as obligate intracellular parasites, viruses have evolved numerous mechanisms to prevent and antagonize innate detection by host cells. Coronaviruses (CoVs), which are the largest known positive-sense, single-stranded RNA viruses, encode several type I interferon (IFN) antagonists. Many of these antagonists prevent the induction of IFN, while others mediate resistance to the effects of IFN (1-3). Upon secretion from a cell, IFNs bind to cell surface-expressed IFN-α/β receptors (IFNARs) in an autocrine and paracrine manner. Subsequently, an IFN signaling cascade utilizing the Janus kinase and signal transducer and activator of transcription pathway leads to the induction and expression of hundreds of interferon stimulated genes (ISGs) that act to limit or prevent viral replication and spread (4). However, during CoV infection, nonstructural protein (nsp) 1 antagonizes the innate immune response by degrading host mRNAs and suppressing IFN beta (IFN-β) expression (5, 6). The nsp3 of severe acute respiratory syndrome coronavirus (SARS-CoV) prevents IRF3 phosphorylation and NF-ϰ B signaling (7). In addition, SARS-CoV nsp3 encodes deubiquitinating and deISGylating activities (3, 8). CoV viral RNA evades innate detection by pattern recognition receptors (PRRs) such as MDA5, and antiviral effectors, such as IFIT1, through formation of a 5' cap-1 structure by encoding N7-methyltransferase and 2'O-methyltransferase activities within nsp14 and nsp16, respectively (9, 10). Murine hepatitis virus (MHV) and Middle East respiratory syndrome coronavirus (MERS-CoV) also encode 2'-5' phosphodiesterases that degrade 2'-5' oligoadenylates, which are key signaling molecules generated by oligoadenylate synthetase (OAS) in response to innate detection of dsRNA that subsequently activate RNase L (11). Most recently, a CoV nsp15 endonuclease activity (EndoU) mutant virus was shown to have increased dsRNA levels, suggesting that nsp15 EndoU reduces dsRNA levels during infection (12).

CoV nsp14 encodes 3'-to-5' exoribonuclease (ExoN) and N7-methyltransferase (N7-MTase) activities (9, 13). CoV nsp14 N7-MTase activity is essential for efficient translation of the viral genome and preventing innate detection (14). In addition, initial biochemical studies of nsp14 ExoN activity demonstrated that ExoN has a preference for dsRNA and the capacity to excise 3' end misincorporated nucleotides (13). Moreover, nsp14 ExoN activity is required for high-fidelity replication. The CoV nsp14 ExoN is a member of the DE-D-Dh superfamily of DNA and RNA exonucleases, so named for the three motifs of four active site residues (13). *Betacoronaviruses* SARS-CoV and MHV expressing engineered, ExoN-inactivating substitutions at active site residues in Motif I (DE → AA) [ExoN(-)] demonstrate increased mutation frequencies and are profoundly sensitive to inhibition by RNA mutagens (15, 16). Additionally, SARS-CoV ExoN(-) virus is attenuated *in vivo* (17). Interestingly, outside of the order *Nidovirales*, the only other known RNA virus-encoded 3'-to-5' exoribonucleases are found in the *Arenaviridae* family of viruses. Lassa fever virus nucleoprotein ExoN is not thought to participate in fidelity regulation, but rather it participates in immune evasion by degrading dsRNA and thereby prevents antigen-presenting cell-mediated NK cell activation (18-20). Recently, in the *Alphacoronavirus* transmissible gastroenteritis virus (TGEV), a mutation in the nsp14 ExoN zinc finger was shown to generate lower levels of dsRNA compared to wild-type (WT) TGEV. However, in that study viruses with mutations in ExoN active site motifs were non-viable and therefore, could not be directly tested for effects on innate immunity (21).

Here, we demonstrate that viruses lacking ExoN activity were sensitive to the effects of IFN pretreatment. In addition, for viruses lacking ExoN activity, replication was restricted in wild-type bone marrow derived macrophages (B6, BMMs) but restored in interferon alpha/beta receptor deficient (IFNAR-/-) BMMs. Despite an increased sensitivity to the effects of IFN treatment, MHV ExoN mutants failed to induce detectable IFN-β gene expression or RNase L-mediated ribosomal RNA (rRNA) degradation and only a limited decrease in viral RNA accumulation was observed. Finally, ExoN(-) virus replicated in the presence of an IFN-βmediated antiviral state had both a decreased specific infectivity and decreased relative fitness compared to untreated ExoN(-) virus. Thus, nsp14 ExoN appears to block or correct the restriction of MHV infection by an IFN-mediated mechanism that may involve damaging nascent viral RNA and affecting subsequent infectivity.

## RESULTS

### Viruses lacking ExoN activity are sensitive to the effects of IFN-β

Binding of type I interferon to the IFNAR receptor on the cell surface leads to a Jak/STAT signaling cascade that ultimately results in the up-regulation and expression of hundreds of antiviral ISGs (4). In addition, WT-MHV replication has been shown to be relatively resistant to the effects of IFN (1, 3, 22). To determine whether the ExoN activity of MHV nsp14 was required for resistance to IFN, we pretreated murine delayed brain tumor (DBT) cells with increasing concentrations of mouse IFN-β for 18 h prior to infecting with WT-MHV or ExoN(-) virus at a multiplicity of infection (MOI) of 1 plaque-forming unit (PFU) per cell (Fig. 1A). In response to IFN-β pretreatment, WT-MHV viral titer decreased by approximately 1 log_10_ as previously reported (1). In contrast, ExoN(-) viral titer demonstrated a dose-dependent decrease and resulted in an approximately 3 log_10_ decrease in viral titer relative to untreated ExoN(-) viral titers. The ExoN activity of nsp14 is conferred by active site residues present in 3 different motifs within the ExoN domain (23). Therefore, to determine whether the observed sensitivity to IFN-β pretreatment for ExoN(-) virus in Fig. 1A was due specifically to the absence of ExoN activity in nsp14, we engineered and recovered a virus encoding only an aspartic acid to alanine substitution in Motif III [ExoN3 (-)]. Previously, we have demonstrated that viruses lacking ExoN activity have decreased replication fidelity and are sensitive to the RNA mutagen 5-fluorouracil (5-FU) (16). Hence, 5-FU sensitivity is an *in vitro* indicator of ExoN activity. Therefore, first, we tested whether ExoN3 (-) and ExoN(-) demonstrated similar sensitivity to 5-FU to ensure that the ExoN activity of ExoN3 (-) virus had been ablated. Similar to ExoN(-), ExoN3 (-) viral replication in cells treated with increasing concentrations of 5-FU demonstrated a dose-dependent decrease in viral titer relative to vehicle treated cells (Fig. 1B). Further, ExoN(-) and ExoN3(-) displayed similar sensitivities to pre-treatment with 100 or 500 U/mL IFN-β following infection at an MOI of 1 PFU/cell (Fig. 1C). Thus, these data suggest nsp14 ExoN activity is required for resistance to the effects of IFN-β pretreatment and subsequent up-regulation and expression of ISGs.

**FIG 1.**
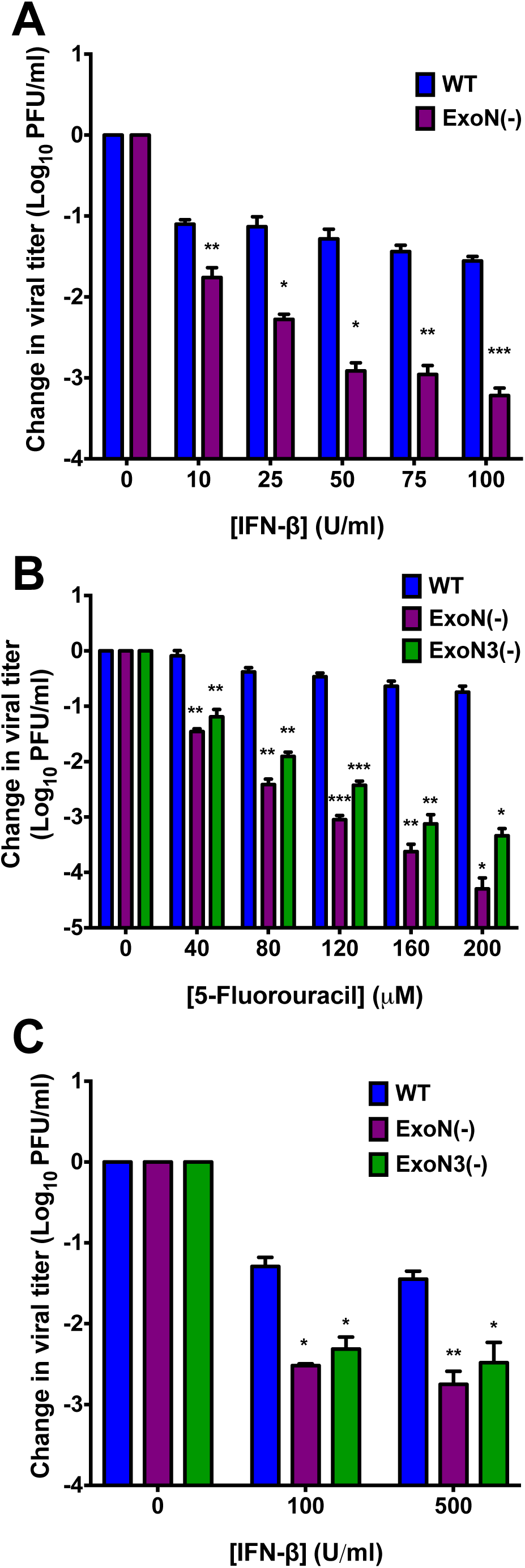
Viruses lacking ExoN activity are sensitive to IFN-β pretreatment. (A) DBT cells were pretreated with the indicated concentrations of mouse IFN-β for 18 h and then infected with WTMHV or ExoN(-) virus (A) or WT, ExoN(-), or ExoN3 (-) virus (C) at an MOI of 1 PFU/cell. At 12 h post-infection, cell culture supernatants were collected and the viral titers present determined by plaque assay. (B) DBT cells were pretreated with the indicated concentrations of 5-FU for 30 min. Following pretreatment, cells were infected with WT, ExoN(-), or ExoN3 (-) virus at an MOI of 1 PFU/cell for 45 min., inocula were removed, and fresh medium containing vehicle or the appropriate concentration of 5-FU were added. Cell culture supernatants were harvested 12 h post-infection and viral titers were determined by plaque assay. For each panel, the change in viral titer was calculated by dividing viral titers following the indicated treatment by the untreated controls and error bars indicate SEM (n = 4). Statistical significance compared to WT-MHV is denoted and was determined by Student's *t* – test. *, *P* < 0.05, \*\**P* < 0.01, *** *P* < 0.001.

### Increased replication capacity does not confer resistance to the effects of IFN-β pretreatment for viruses lacking ExoN activity

ExoN(-) virus demonstrates an approximately 2 h delay in exponential replication and a 1 log_10_ decrease in peak titer relative to WT-MHV (15). Therefore, we tested whether the IFN sensitivity phenotype observed for ExoN(-) and ExoN3 (-) viruses is due to the decreased replication capacity of these viruses. To do so, we utilized an ExoN(-) virus developed by our lab that has been blindly passaged in DBT cells for 250 passages [ExoN(-) P250] (24). The replication capacity of the resulting ExoN(-) P250 virus exceeds that of WT-MHV (Fig. 2A). However, despite increased replication capacity, ExoN(-) P250 demonstrated similar sensitivity to IFN-β pretreatment as ExoN(-) virus (Fig. 2A and B). Hence, the IFN-β sensitivity phenotype of viruses lacking ExoN activity is not dependent on viral replication capacity but instead, is directly associated with a specific function of nsp14 ExoN that is required for efficient replication in the presence of an IFN-β-mediated antiviral state.

**FIG 2.**
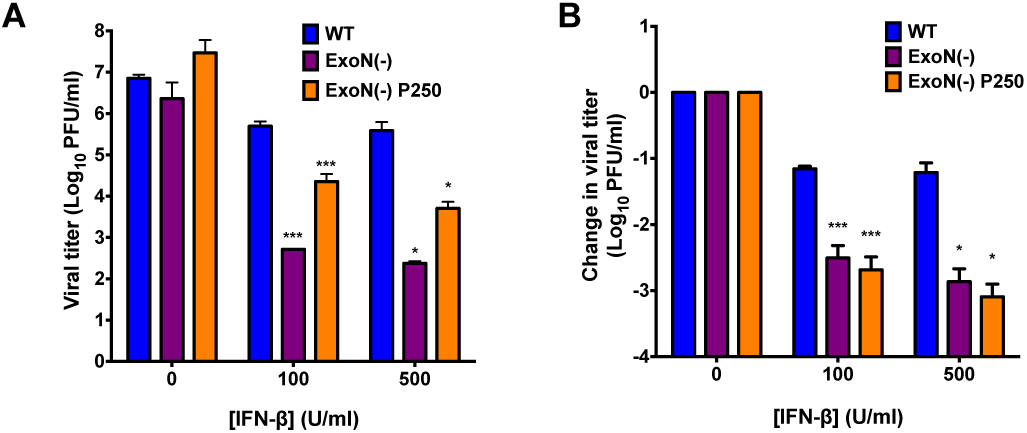
Increased replication capacity does not restore virus resistance to IFN-β. DBT cells were pretreated with the indicated concentrations of mouse IFN-β for 18 h and then infected with WT, ExoN(-), or ExoN(-) P250 virus at an MOI of 1 PFU/cell. At 12 h post-infection, cell culture supernatants were collected and the viral titers present determined by plaque assay. Raw viral titers (A) or the change in viral titers relative to untreated controls (B) are reported. Error bars indicate SEM (n = 4). Statistical significance compared to WT-MHV is denoted and was determined by Student's *t* – test. *, *P* < 0.05, \*\**P* < 0.01, *** *P* < 0.001.

### Nsp14 ExoN activity is required for replication in wild-type B6 BMMs

We next wanted to test whether ExoN activity was required for IFN resistance in primary innate immune cells, such as BMMs. Replication of WT-MHV in primary BMMs is well described, and data suggest that wild-type B6 BMMs (B6) express many PRRs and ISGs at a higher basal level than many mouse cell lines (3, 25, 26). In contrast, BMMs lacking the IFNAR receptor (IFNAR-/-) have lower basal and expressed levels of ISGs; thus, making B6 and IFNAR-/-BMMs excellent cell types for interrogating the role of ExoN activity on viral replication and antagonism of the innate immune response (25). BMMs from B6 or IFNAR-/-mice were generated and infected with WT-MHV or ExoN(-) virus at an MOI of 1 PFU/cell. Samples were harvested at the indicated time points, and viral titers were determined by plaque assay (Fig. 3A). WT-MHV replication increased gradually in both B6 and IFNAR-/-BMMs at each time-point post-infection. In contrast, ExoN(-) virus replication in B6 BMMs was only detectable at 6 and 9 h post-infection. However, when IFNAR-/- BMMs were infected with ExoN(-) virus, viral titers were partially restored and increased at each time point post-infection. To further test the replication of viruses lacking ExoN activity and the effect of an increased replication capacity in BMMs, B6 and IFNAR-/-BMMs were infected with WT-MHV or ExoN(-) P250 viruses at an MOI of 1 PFU/cell. Similar to Fig. 3A, WT-MHV viral titers steadily increased in B6 and IFNAR-/-BMMs at each time-point post-infection (Fig. 3B). However, similar to ExoN(-) virus, ExoN(-) P250 virus replication in B6 BMMs was restricted and not detected beyond 9 h post-infection. In addition, ExoN(-) P250 virus replication in IFNAR-/-BMMs was restored to similar levels as WT-MHV. These data show that ExoN activity is required for replication in B6 BMMs. Further, they suggest that restriction of ExoN(-) or ExoN(-) P250 is mediated by a gene or genes downstream of IFNAR.

**FIG 3.**
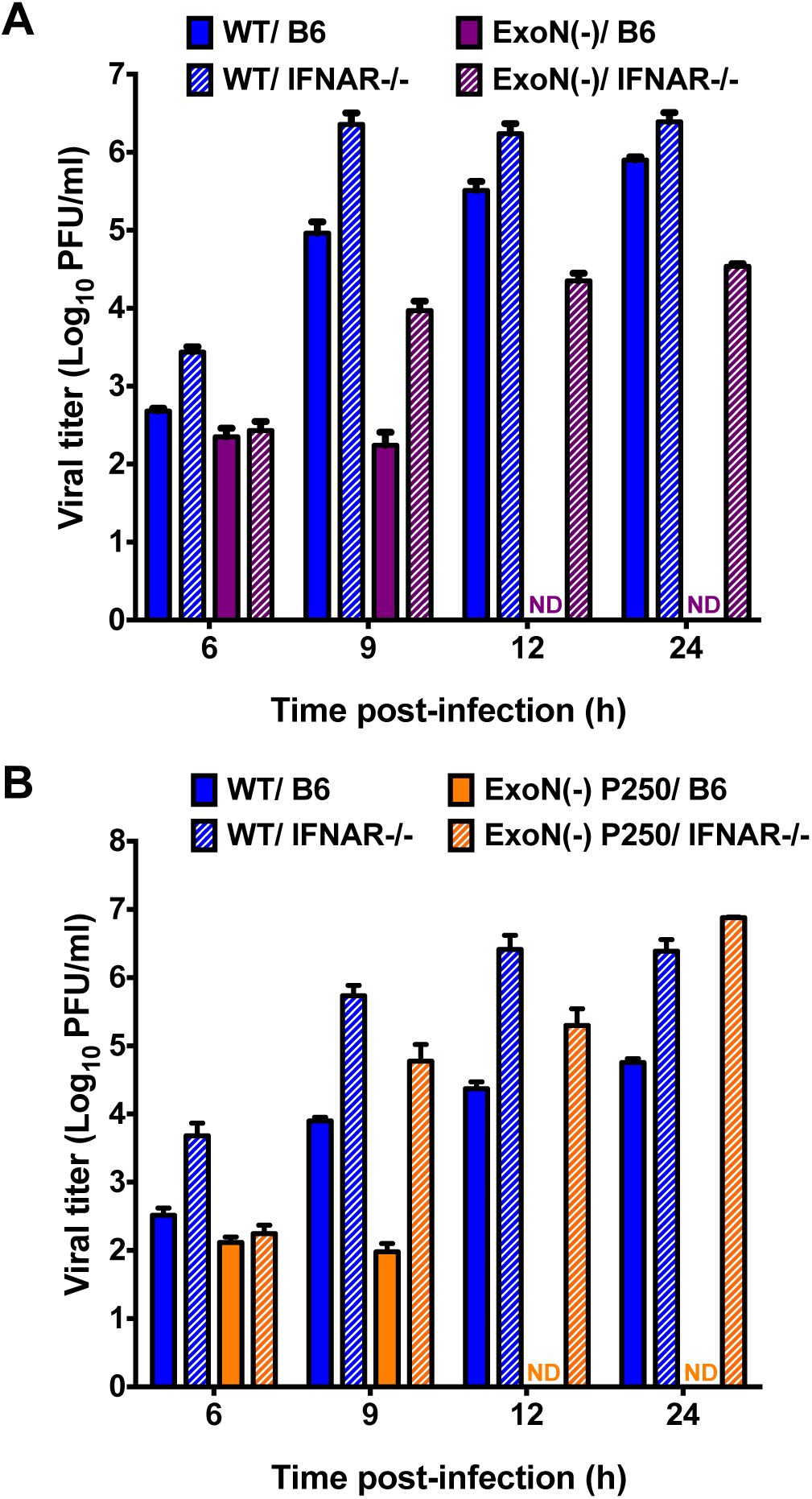
Replication of viruses lacking ExoN activity is restricted in wild-type B6 BMMs. B6 BMMs or IFNAR-/-BMMs were infected with WT-MHV or ExoN(-) virus (A) or WT-MHV or ExoN(-) P250 virus (B) at an MOI of 1 PFU/cell. At the indicated times post-infection, cell culture supernatant aliquots were collected and the viral titers present were determined by plaque assay. For each panel, error bars represent SEM (n = 6 to 7). ND = not detectable.

### Loss of ExoN activity does not result in the induction of IFN and replication is not rescued by RNase L/PKR deficiency

Upon detection of a pathogen-associated molecular pattern (PAMP) by innate sensors, signaling pathways lead to transcription factor activation and nuclear translocation resulting in expression of IFN-β mRNA (4). WT-MHV is well known to prevent or delay the induction of IFN expression (3, 22). However, ExoN activity may help prevent the detection of a PAMP, namely dsRNA, which has been shown to be increased in an nsp15 EndoU mutant (12). Therefore, to determine whether the loss of ExoN activity resulted in the generation and subsequent detection of a PAMP, we determined the level of IFN-β gene expression in DBT cells infected with mock, WT-MHV, ExoN(-), or ExoN(-) P250 virus at an MOI of 0.1 PFU/cell (Fig. 4A). In addition, we infected DBT cells with Sendai virus (SenV), a positive control and a potent inducer of IFN, at an MOI of 200 HA units/ml. SenV infection resulted in IFN expression by 3 h post-infection and peaked between 9 and 12 h post-infection prior to returning nearly to mock infected levels by 24 h post-infection, demonstrating that DBT cells are capable of expressing IFN-β. In contrast, no CoV infection, regardless of whether intact ExoN activity was present, resulted in IFN-β gene expression over mock-infected cells with the exception of WT-MHV at 3 h post-infection. Further, upon detection of dsRNA by OAS and subsequent activation of RNase L, viral and cellular RNAs are degraded as an antiviral mechanism (4). To determine whether infection with ExoN(-) virus activates RNase L, DBT cells were pretreated with 0 or 50 U/ml mouse IFN-β and infected with WT-MHV or ExoN(-) virus at an MOI of 1 PFU/cell or transfected with 25μg/ml poly I:C, a dsRNA surrogate. At the indicated times post-infection, cell lysates were harvested, total RNA extracted, and the integrity of cellular rRNA determined using a bioanalyzer (Fig. 4B). Transfection of DBT cells with poly I:C resulted in rRNA degradation, whereas, infection of DBT cells with WT-MHV or ExoN(-) virus did not result in rRNA degradation under any tested conditions. Lastly, when B6 or RNase L-/-/PKR-/-BMMs (RL-/-/PKR-/-) were infected with ExoN(-) virus, replication was restricted (Fig. 4C). In contrast to infection of B6 BMMs, ExoN(-) viral titer from RL-/-/PKR-/-BMMs was detectable at 12 and 24 h post-infection. However, viral yield was minimal. These data suggest that loss of nsp14 ExoN activity does not lead to the transcriptional activation of IFN-β or a notable dsRNA sensor such as OAS/RNase L during infection of DBT cells. In addition, BMMs deficient in the antiviral effectors RNase L and PKR were not sufficient to restore ExoN(-) viral replication.

**FIG 4.**
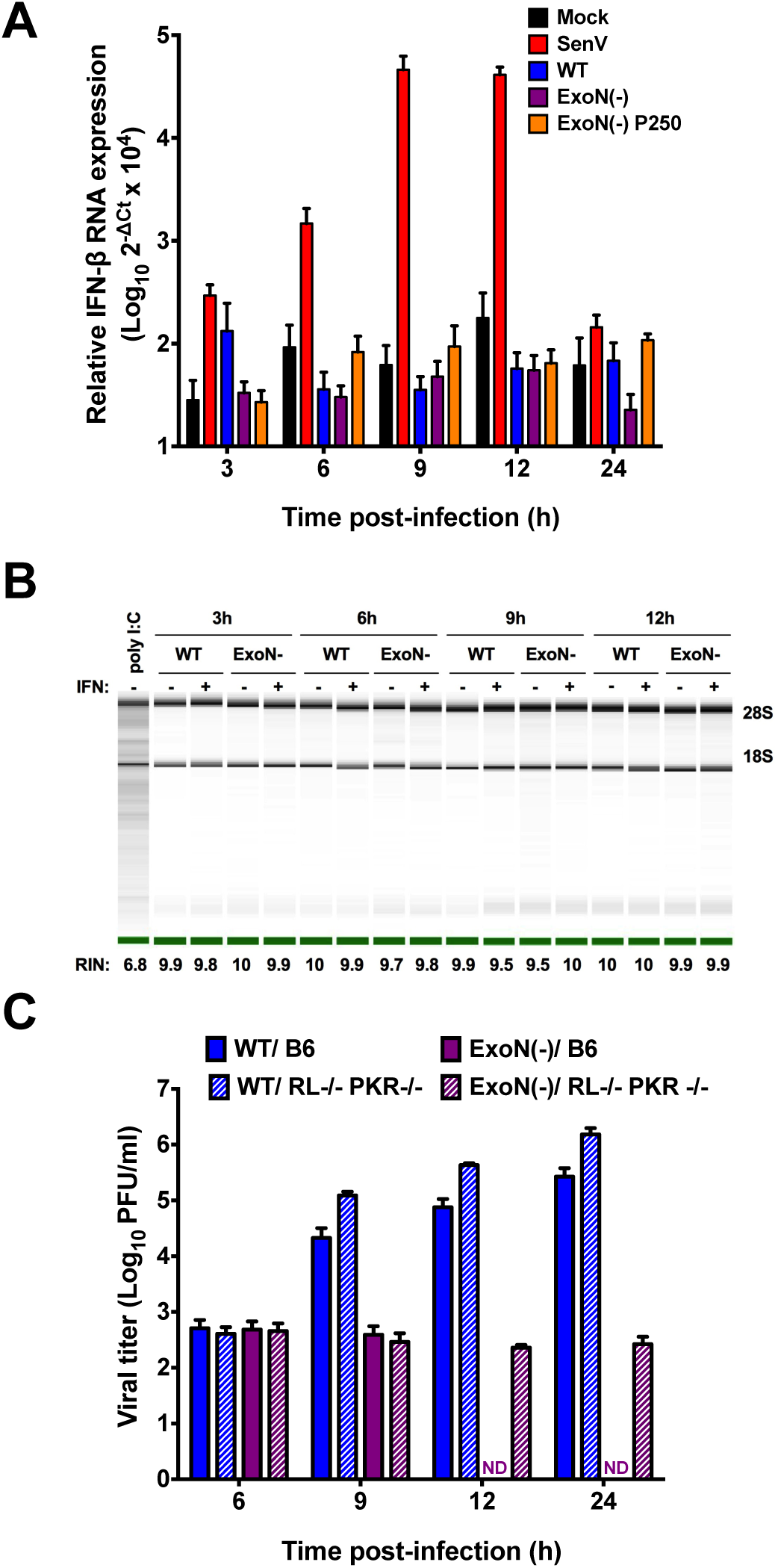
Loss of ExoN activity does not result in the generation of a detectable PAMP. (A) DBT cells were infected with mock, WT, ExoN(-) or ExoN(-) P250 virus at an MOI of 0.1 PFU/cell or infected with Sendai virus at an MOI of 200 HA units/ml. At the indicated times post-infection, cell culture supernatants were removed, cell lysates were harvested, total RNA was extracted, cDNA was generated, and IFN-β expression relative to GAPDH was determined by qPCR. Error bars indicate SEM (n = 4). (B) DBT cells were pretreated for 18 h with 0 or 50U/ml mouse IFN-β and subsequently infected with WT-MHV or ExoN(-) virus or transfected with 25μg/ml poly I:C. At the indicated times post-infection, cell culture supernatants were removed, cell lysates harvested, and total RNA extracted. rRNA integrity was assessed using an Agilent Bioanlyzer. One representative image is shown for each sample from 2 independent experiments. Images spliced for labeling purposes. The averaged RNA integrity values for each condition are reported. (C) B6 BMMs or RL-/-/ PKR-/-BMMs were infected with WT-MHV or ExoN(-) virus at an MOI of 1 PFU/cell. At the indicated times post-infection, cell culture supernatant aliquots were collected and the viral titers present were determined by plaque assay. Error bars represent SEM (n = 5). ND = not detectable.

### IFN treatment does not substantially alter ExoN(-) viral RNA accumulation or particle release

Since ExoN activity is required for resistance to IFN but had no effect on IFN induction, we sought to discern the stage of viral replication that was restricted by IFN-β treatment. To determine the effect of IFN-β pretreatment on viral RNA accumulation, DBT cells were pretreated with 0 or 100 U/ml mouse IFN-β for 18 h and subsequently infected with WT-MHV or ExoN(-) virus at an MOI of 1 PFU/cell. At the indicated times post-infection, the number of genomic RNA copies present were determined by qRT-PCR. IFN-β pretreatment had minimal effect on the accumulation of WT-MHV genomic RNA (Fig. 5A). Whereas ExoN(-) genomic RNA accumulation is delayed relative to WT-MHV (15), pretreatment with IFN-β did not substantially decrease ExoN(-) genomic RNA levels (Fig. 5A). In addition, we determined the effects of IFN-β pretreatment on the levels of subgenomic viral RNA. For both WT-MHV and ExoN(-) viruses, IFN-β pretreatment did not substantially reduce subgenomic RNA levels at any time-point (Fig. 5B). These data indicate that IFN pretreatment did not result in the gross degradation or inhibition of ExoN(-) or ExoN(-) P250 viral RNA accumulation. While slight reductions in viral RNA could explain a small portion of the IFN phenotype, these data suggest that decreased replication or transcription is not the primary driver of ExoN(-) IFN sensitivity.

**FIG 5.**
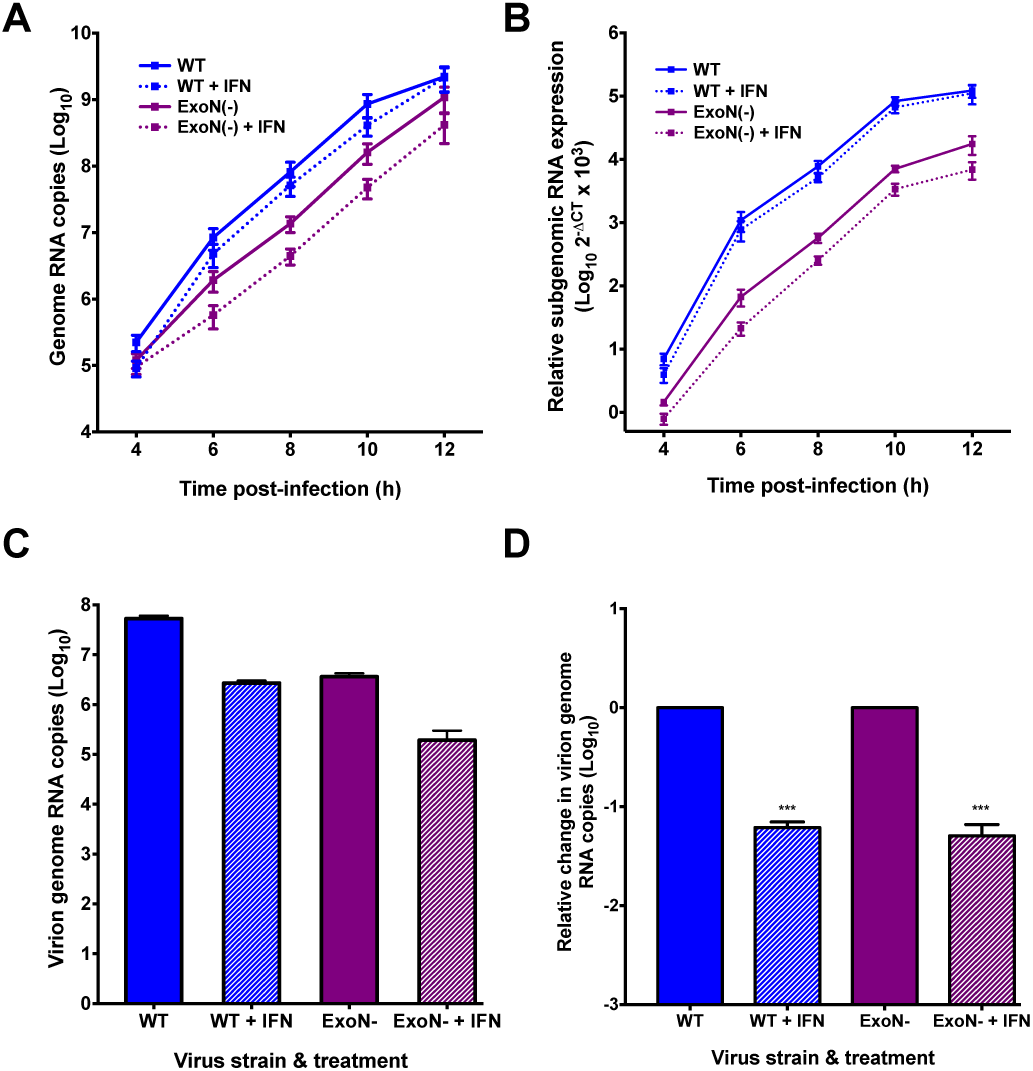
ExoN(-) viral RNA accumulation and particle release is marginally affected by IFN-β pretreatment. DBT cells were pretreated with 0 or 100U/ml mouse IFN-β for 18 h and subsequently infected with WT-MHV or ExoN(-) virus at an MOI of 1 PFU/cell. At the indicated times post-infection, total cell lysates were harvested and RNA was extracted. The viral genomic RNA copies present relative to an RNA standard were determined by one-step qRT-PCR (A) or cDNA was generated and the subgenomic RNA copies relative to GAPDH were determined by qPCR (B). For each panel (A and B), error bars represent SEM (n= 6 to 9). DBT cells were pretreated with 0 or 100U/ml mouse IFN-β for 18 h and subsequently infected with WT-MHV or ExoN(-) virus at an MOI of 1 PFU/cell. At 12 h post-infection, cell culture supernatants were collected. Equivalent volumes of cell culture supernatant for each sample were divided into two samples. For the first cell culture supernatant sample, total RNA was extracted and the number of virion genome RNA copies present (particles) was determined by one-step qRT-PCR (C) or reported as the change in virion genome RNA copies (D). Error bars represent SEM (n = 13 to 15). Statistical significance compared to untreated WT-MHV or ExoN(-) infection, respectively, is denoted and was determined by Student's *t* – test. *** *P* < 0.001.

Since pretreatment of DBT cells with IFN-β does not grossly alter ExoN(-) viral RNA accumulation but does reduce ExoN(-) viral titers, we sought to determine whether IFN pretreatment prior to infection resulted in a measurable difference in the number of viral particles released from WT-MHV or ExoN(-) infected cells. DBT cells were pretreated with 0 or 100 U/ml mouse IFN-β for 18 h and subsequently infected with WT-MHV or ExoN(-) virus at an MOI of 1 PFU/cell. At 12 h post-infection, cell culture supernatants were harvested and an aliquot of two equal volumes were removed. From the first volume of each sample, RNA was extracted and used to perform one-step qRT-PCR to determine the number of genome RNAs present, and hence, the number of genome RNA containing particles present in the given volume of supernatant (Fig. 5C). The second volume was saved for a plaque assay as described below. Pretreatment of cells with IFN-β resulted in approximately a 1 log_10_ decrease in the number of supernatant viral particles for both WT-MHV and ExoN(-) viruses compared to the number of supernatant viral particles from untreated cells, demonstrating that IFN pretreatment affects the release of WT-MHV and ExoN(-) virus particles equally (Fig. 5D). Thus, these data suggest that IFN pretreatment does not restrict the primary replication of viruses lacking ExoN activity, but rather, renders them potentially inadequate for subsequent infection.

### ExoN(-) virus progeny generated in the presence of an IFN induced-antiviral state have decreased specific infectivity and fitness upon subsequent infection

While many ISGs antagonize viral replication, some could alter the infectivity of progeny particles (27, 28). To test whether IFN pretreatment affected the infectivity of ExoN(-) viral particles, the remaining cell culture supernatant volume described above was used to perform a plaque assay to determine the number of PFUs present (data not shown). Using the number of particles determined in Fig. 5C and the number of PFUs present in an equivalent volume; we calculated the specific infectivity, or particle-to-PFU ratio, of each virus generated under each condition (Fig. 6A). Regardless of IFN-β pretreatment during the initial infection, the specific infectivity of WT-MHV was approximately 10 particles per 1 PFU upon subsequent infection. Infection of untreated DBT cells with ExoN(-) virus resulted in a similar particle to PFU ratio as WT-MHV during subsequent infection. In contrast, when DBT cells were pretreated with IFN-β prior to initial infection with ExoN(-) virus, the resulting specific infectivity of ExoN(-) virus was 100 particles per 1 PFU, a significant decrease in specific infectivity. Therefore, ExoN(-) virus generated in the presence of an IFN-β-mediated antiviral state requires 10-fold more genome RNA containing particles to generate 1 PFU than WT-MHV generated in cells pretreated with or without IFN or ExoN(-) virus generated in untreated cells. These data suggest that the IFN-mediated restriction of ExoN(-) virus in DBT cells occurs at the level of subsequent infection by reducing particle infectivity.

**FIG 6.**
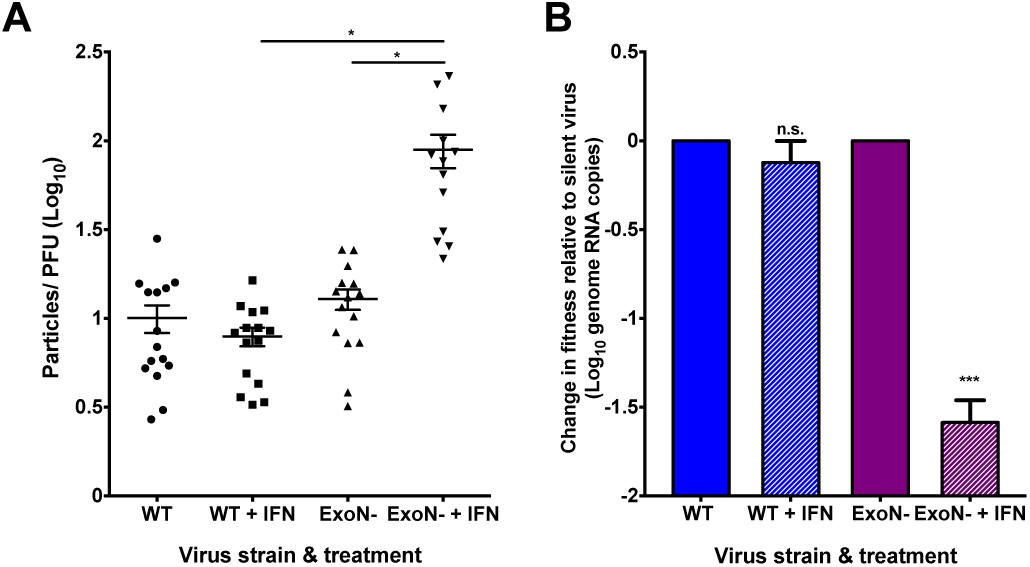
ExoN(-) viruses generated in the presence of an antiviral state have decreased specific infectivity and are less fit relative to untreated. (A) DBT cells were pretreated with 0 or 100U/ml mouse IFN-β for 18 h and subsequently infected with WT-MHV or ExoN(-) virus at an MOI of 1 PFU/cell. At 12 h post-infection, cell culture supernatants were collected. Equivalent volumes of cell culture supernatant for each sample were divided into two samples. For the first cell culture supernatant sample, total RNA was extracted and the number of virion genome RNA copies present (particles) was determined by one-step qRT-PCR [Fig. 5C]. For the second cell culture supernatant sample, the viral titer present was determined by plaque assay (PFUs) (data not shown). The particle to PFU ratio for each virus and treatment was calculated by dividing the number of particles by the number of PFUs. Error bars represent SEM (n = 13 to 15). (B) DBT cells were pretreated with 0 or 100 U/ml mouse IFN-β for 18 h and subsequently infected with WT-MHV or ExoN(-) virus at an MOI of 1 PFU/cell. At 12 h post-infection, cell culture supernatants were harvested for each virus and treatment group and the number of virion genome RNA copies present (particles) in the supernatant was determined by one-step qRT-PCR. Using the determined number of particles, an equivalent number of virus particles from each virus and treatment group were mixed with an equal number of WT silent or ExoN(-) silent virus particles. This mixture was then used to infect a fresh monolayer of untreated DBT cells. At 24 h post-infection, cell culture supernatants were collected, RNA was extracted, and the number of virion genome RNA copies for each original virus and treatment group relative to their respective silent standard viruses was determined by one-step qRT-PCR and is reported as the change in fitness relative to the silent virus standard. Error bars represent SEM (n = 6). For each panel, statistical significance compared to untreated WT-MHV or ExoN(-) infection, respectively, is denoted and was determined by Student's *t* – test. *, *P* < 0.05, *** *P* < 0.001, n.s. = not significant.

Next, we tested whether the effects of IFN on ExoN(-) viruses were intrinsic to the viruses produced. To do so, we performed a co-infection assay, which utilized WT-MHV and ExoN(-) viruses harboring 10 silent mutations in the nsp2-coding region (WT silent and ExoN(-) silent, respectively) along with WT-MHV and ExoN(-) viruses. The genome RNAs of the silent viruses are recognized exclusively by a separate probe than the one used to detect the “WT” nsp2 probe-binding region of WT-MHV or ExoN(-) viruses (24). Thus, allowing WT silent and ExoN(-) silent to act as internal controls for a co-infection assay under identical conditions. DBT cells were pretreated with 0 or 100 U/ml mouse IFN-β for 18 h and subsequently infected with WT-MHV or ExoN(-) virus at an MOI of 1 PFU/cell. At 12 h post-infection, total cell culture supernatants were collected. The number of viral particles present in a representative aliquot was determined from purified virion genome RNA by one-step qRT-PCR. In addition, we determined the number of genome RNA containing particles in an equivalent volume of WT silent or ExoN(-) silent viral p1 stock tubes. Using the calculated number of genome RNA-containing viral particles, we added an equal number of WT-MHV viral particles generated in the absence of IFN pretreatment to WT silent viral particles and an equal number of WT-MHV viral particles generated in the presence of IFN pretreatment to WT silent viral particles. This same set-up was repeated for ExoN(-) viral particles generated in the presence or absence of IFN pretreatment with ExoN(-) silent viral particles. Finally, each combination was used to infect a fresh monolayer of untreated DBT cells. At 24 h post-co-infection, total cell culture supernatants were collected and virion genome RNA was extracted to determine the number of supernatant viral particles present from each combination of input viruses by one-step qRT-PCR and is reported as the change in fitness relative to the respective silent virus standard (Fig. 6B). WT-MHV particles generated in the presence of IFN pretreatment generated a similar number of viral particles over the course of co-infection as WT-MHV generated in the absence of IFN pretreatment relative to their respective silent standards. However, the number of viral particles present from ExoN(-) virus generated in the presence of IFN pretreatment during the course of co-infection decreased by approximately 1.5 log_10_ in comparison with ExoN(-) viral particles generated in the absence of IFN pretreatment relative to their respective silent standards. These data indicate that a loss in nsp14 ExoN activity sensitizes viruses to IFN pretreatment and reduces the infectivity and fitness of progeny during subsequent rounds of infection in the absence of an antiviral state.

## DISCUSSION

CoVs encode multiple IFN antagonists that prevent the induction of or mediate resistance to the innate immune response; thus, allowing efficient viral replication early during infection (3). Moreover, an insufficient innate immune response has been proposed to be a major contributor to SARS-CoV pathogenesis (29). In this study, we sought to determine the contributions of nsp14 ExoN activity in the induction of and resistance to the innate immune response. We demonstrate that ExoN(-) virus is sensitive to pretreatment with IFN-β. Because ExoN3 (-) and ExoN(-) P250 viruses were also sensitive to the effects of IFN, we conclude that IFN sensitivity is specifically due to loss of ExoN activity.

Because the ExoN activity of the Lassa fever virus nucleoprotein degrades dsRNA intermediates (18, 19), we hypothesized that CoV nsp14 ExoN could function in a similar manner. If nsp14 ExoN is degrading viral dsRNA, ExoN inactivation should increase intracellular dsRNA accumulation, resulting in a concomitant increase in IFN-β expression or activation of RNase L during infection. We neither observed IFN-β up-regulation nor RNase L activation over the course of ExoN(-) virus infection (Fig. 4A and B), and rRNA was intact at all time-points tested (Fig. 4B). Therefore, at least two possible explanations exist: 1.) ExoN does not function to degrade dsRNA or 2.) ExoN does degrade dsRNA, but the detection of this PAMP is unchanged during ExoN(-) virus infection due to sufficient antagonism by other CoV proteins. Basal OAS expression levels correlate with RNase L activation (30). Thus, we pretreated DBTs with IFN-β to up-regulate OAS and RNase L expression. However, rRNA degradation was only observed in cells transfected with poly I:C (Fig. 4B). Further, nsp15 EndoU and NS2 phosphodiesterase activities were intact during all of our experiments. Thus, it is possible that in the absence of nsp14 ExoN activity, other CoV innate antagonists were sufficient to prevent innate detection by the cell or prevent the induction of a detectable signal in the experiments we performed. However, one would expect the endonucleolytic products of nsp15 to be smaller dsRNAs that could still activate RIG-I or MDA5, similar to RNase L products, unless another RNA degradation mechanism were in place (4, 31). In addition, despite an intact NS2 phosphodiesterase, nsp15 mutants still activate RNase L-mediated rRNA degradation (12). Lastly, when RNaseL-/-/ PKR-/-BMMs were infected with ExoN(-) virus, viral replication was not rescued, suggesting that RNase L and PKR are not required for ExoN(-) virus restriction (Fig. 4C). Moreover, these data suggest dsRNA is not detected and the antiviral effectors RNaseL and PKR are not activated during ExoN(-) virus infection.

During our study, Becares et al. reported that a TGEV nsp14 zinc-finger mutant modulated the innate immune response of swine testis cells by reducing the levels of dsRNA and induction of IFN (21). Unlike TGEV, *Betacoronaviruses* such as SARS-CoV and MHV, do not induce IFN expression in most cell types (1-3) (Fig.4A). Interestingly, TGEV ExoN active site mutants were non-viable; although, this is not the first report of non-viable ExoN active site residue mutants in *Alphacoronaviruses* (21). In the initial report of CoV nsp14 ExoN activity, human CoV 229E ExoN active site mutants were also non-viable, suggesting a common essential function for nsp14 ExoN in *Alphacoronavirus* replication and/or innate antagonism (13). Altogether, the possibility of a common innate immune antagonism function for nsp14 across *Alpha*-and *Beta*-CoVs is apparent but clearly differing requirements exist that may be dependent on the CoV genus and cell types used.

Our results clearly demonstrate that viruses lacking ExoN activity are sensitive to IFN-β pretreatment in a dose-dependent manner (Fig. 1A, C and Fig. 2). Further, replication of viruses lacking ExoN activity was dependent on the capacity of BMMs to express genes downstream of IFNAR signaling (Fig. 3). This is due to the fact that B6 and IFNAR-/-cells have different levels of basal ISG expression and thus, two very different intracellular environments for viral replication to occur (3, 25, 26). In IFNAR-/-BMMs, ExoN(-) and ExoN(-) P250 virus replication capacity was restored to levels approaching or exceeding WT-MHV levels (Fig. 3). Further, our specific infectivity (Fig. 6A) and co-infection (Fig. 6B) data show that ExoN(-) virus generated in the presence of an antiviral state is less viable upon subsequent infection. Altogether, our results suggest that an ISG or ISGs is (are) acting on ExoN(-) virus, specifically resulting in progeny that are less viable upon subsequent infection. Thus, it will be interesting to determine the specific ISG or ISGs responsible for mediating the observed restriction. In addition, it will be important to determine whether a greater proportion of the incoming ExoN(-) viral particles are strictly non-viable or whether cells are now sensing the progeny ExoN(-) viruses and inhibiting replication. Due to the pleiotropic nature of IFN-β, more than one mechanism may be acting. To date, the majority of our understanding of nsp14 ExoN activity is in the context of proofreading during CoV replication (13, 15, 16, 32). CoVs lacking ExoN activity demonstrate an increase in mutation frequency relative to WT (15, 16). Thus, it is possible that ExoN(-) virus replication in IFN pretreated cells results in further alteration of ExoN(-) virus mutation frequency. Certainly, an increase or decrease in mutation frequency could impair viral replication during a subsequent infection. In addition, an ISG may act to hypermutate the large CoV genome in the absence of ExoN activity, rendering viral progeny less viable. ISGs that increase viral mutation frequency have been described such as adenosine deaminase acting on RNA 1 (ADAR1) and apolipoprotein B mRNA editing enzyme, catalytic polypeptide-like 3G (APOBEC3G) (27, 28). Further, another ISG, SAMHD1, may inhibit HIV replication by limiting nucleotide pools, a known contributor to increased viral mutation frequency (33-35). Moreover, other possible mechanisms outside of altered mutation frequency exist. For instance, in the absence of ExoN activity, terminal RNA modifications, recombination, and/or replicase protein interactions mediated by nsp14 ExoN may be disrupted to a greater extent in the presence of an IFN-β-mediated antiviral state.

Since CoVs encode the largest genome known for RNA viruses, they have the luxury of encoding multiple IFN antagonists that limit the capacity of a cell to detect and respond to infection. Collectively, our data suggest that MHV nsp14 ExoN activity is a contributor to CoV innate immune antagonism. We clearly demonstrate that viruses lacking ExoN activity are sensitive to the effects of an IFN-β-mediated antiviral state. Further, our data reveal a critical role for nsp14 ExoN activity in CoV replication and provide additional rationale for targeting nsp14 ExoN activity as a means of viral attenuation. Our future studies will probe the specific mechanism of restriction for viruses lacking ExoN activity and assess how the requirement of ExoN activity for resistance to innate immunity can be utilized for treatment during human coronavirus infections.

## MATERIALS AND METHODS

### Cell culture

Murine delayed brain tumor (DBT) cells (36) and baby hamster kidney 21 cells expressing the MHV receptor (BHK-R) (37) were maintained at 37°C in Dulbecco's modified Eagle medium (DMEM; Gibco) supplemented with 10% fetal bovine serum (FBS; Invitrogen), 100 U/ml penicillin and streptomycin (Gibco), and 0.25 μg/ml amphotericin B (Corning). BHKR cells were further supplemented with 0.8 mg/ml of G418 (Mediatech).

### Cloning, recovery, and verification of mutant viruses

Recombinant MHV strain A59 (GenBank accession number AY910861) has been previously described (37). ExoN(-) (nsp14 D89A and E91A) has been previously described (15). To generate ExoN(-) P250 virus, sub-confluent monolayers of DBT cells in 25cm^2^ flasks were infected using the ExoN(-) parental stock and blindly passaged for a total of 250 passages (24). For ExoN3 (-) virus (nsp14 D272A), site-directed mutagenesis was used to engineer point mutations in the MHV genome cDNA F fragment plasmid using the MHV infectious clone reverse genetics system (37). ExoN3 (-) mutant virus was recovered using BHK-R cells following electroporation of *in vitro*-transcribed genomic RNA. Recovered ExoN3 (-) virus was sequenced (GenHunter Corporation, Nashville, TN) to verify the engineered mutations were present and to ensure that no additional mutations were introduced.

### Interferon-β sensitivity assays

Sub-confluent DBT cells were treated for 18 h with the indicated concentrations of mouse IFN-β (PBL Assay Science) prior to infection with virus at a multiplicity of infection (MOI) of 1 plaque-forming unit (PFU) per cell at 37°C for 45 min. After incubation, inocula were removed, cells were washed with PBS, and fresh medium was added. Cell culture supernatants were collected at 12 h post-infection, and viral titers were determined by plaque assay (15).

### 5-FU sensitivity assays

Sub-confluent DBT cells were treated with DMEM supplemented to contain the indicated concentrations of 5-fluorouracil [(5-FU), Sigma] or DMSO alone at 37**°**C for 30 min. After incubation, drug was removed and cells were infected with virus at an MOI of 1 PFU/cell at 37**°**C for 1 h. Inocula were removed, and cells were incubated in medium containing 5-FU or DMSO. Cell culture supernatants were collected at 12 h post-infection, and viral titers were determined by plaque assay.

### Virus replication kinetics

Bone-marrow derived macrophages (BMMs) were generated from the hind limbs of WT, IFNAR-/-, or RNaseL-/-/ PKR-/-C57/B6 mice as previously described (11). BMMs were infected with virus at an MOI of 1 PFU/cell at 37°C for 1 h. After incubation, inocula were removed, cells were washed with 3 times with PBS, and fresh medium was added. At the indicated times post-infection, cell culture supernatant aliquots were collected and viral titers determined by plaque assay.

### Interferon-β induction assays

Sub-confluent DBT cells were infected with mock, WT, ExoN(), or ExoN(-) P250 virus at an MOI of 0.1 PFU/cell or with Sendai virus (SenV) at an MOI of 200 HA (hemagglutination units)/ml at 37°C for 45 min. Inocula were removed, cells were washed with PBS, and fresh medium was added. At the indicated times post-infection, cell culture supernatants were removed and cell lysates were harvested by adding 1ml TRIzol reagent. Total RNA present in the lysates was purified using the phenol/chloroform method. cDNA was generated by reverse transcriptase-polymerase chain reaction (RT-PCR) using 1μg of total RNA as previously described (16). Mouse IFN-β expression levels were determined relative to GAPDH by qPCR using the Applied Biosciences 7500 Real-Time PCR System with Power SYBR Green PCR Master Mix and IFN-β primers: FWD: 5'-TCCGCCCTGTAGGTGAGGTTGAT-3' and REV: 5'-GTTCCTGCTGTGCTTCTCCACCA-3' and GAPDH primers previously reported (16).

### Determination of rRNA integrity

Sub-confluent monolayers of DBT cells were treated with 0 or 50 U/ml mouse IFN-β for 18 h prior to being infected with virus at an MOI of 1 PFU/cell at 37°C for 45 min. After incubation, inocula were removed, cells were washed with PBS, and fresh medium was added. At the indicated times post-infection, cell culture supernatants were removed and total RNA was harvested by adding 1ml TRIzol reagent. For a positive control, cells were transfected with 25ug/ml polyI:C (Sigma) using Lipofectamine 2000 (Thermo Fisher Scientific). Total RNA from all samples was purified using the Purelink RNA Mini Purification System (Life Technologies) by following the manufacturers instructions. Upon purification, total RNA was analyzed on an Agilent Bioanalyzer by the Vanderbilt VANTAGE core facility and the rRNA integrity reported.

### Quantification of viral genomic RNA by qRT-PCR

The quantification of viral genomic RNA has been previously described (14). Briefly, an RNA standard was prepared using the MHV A fragment (37) and a standard curve was generated using 10-fold dilutions from 10^3^ to 10^8^ copies. A 5' 6-carboxyfluorescein (FAM)-labeled probe (5'-TTCTGACAACGGCTACACCCAACG-3' [Biosearch Technologies]) was used with forward (5'-AGAAGGTTACTGGCAACTG-3') and reverse (5'-TGTCCACGGCTAAATCAAAC-3') nsp2 specific primers. The final volume for each reaction was 20 μl with 150 nM probe, 900 nM each primer, 2 μl sample RNA, and 10 μl 2X ToughMix, one-step, low ROX enzyme mix (Quantas) per reaction. Samples were quantified using an Applied Biosciences 7500 Real-Time PCR System with the conditions 55°C for 10 min, 95°C for 5 min, 95°C for 30 s, and 60°C for 1 min, with the last two steps repeated 40 times. The standard curve was plotted using GraphPad Prism 6 software, and genomes/μl were calculated.

### Quantification of subgenomic RNA by qPCR

Sub-confluent DBT cells were treated with 0 or 100 U/mL mouse IFN-β for 18 h prior to being infected with virus at an MOI of 1 PFU/cell at 37°C for 45 min. After incubation, inocula were removed, cells were washed with PBS, and fresh medium was added. At the indicated times post-infection, cell culture supernatants were removed and total RNA was harvested by adding 1ml TRIzol reagent. Total RNA was extracted using the Purelink RNA mini purification system by following the manufacturers instructions. cDNA was generated by RT-PCR using 1ug of total RNA as previously described (16). Primers used to detect subgenomic nucleocapsid and GAPDH gene expression have been reported (16, 38). Subgenomic (N) expression levels relative to GAPDH were determined using the Applied Biosciences 7500 Real-Time PCR System with Power SYBR Green PCR Master Mix.

### Determination of specific infectivity

Sub-confluent monolayers of DBT cells were infected with virus at an MOI of 1 PFU/cell at 37°C for 45 min. After incubation, inocula were removed, cells were washed with PBS, and fresh medium was added. At 12 h post-infection, cell culture supernatants were collected, and viral titers were determined by plaque assay. Supernatants also were used for RNA genome isolation by adding 100 μl supernatant to 900 μl TRIzol reagent, chloroform extraction by phase separation, and final purification using the PureLink RNA Mini Purification System. Genome RNA was quantified using one-step qRT-PCR as described above, and the particle to PFU ratio was calculated.

### Co-infection assay

Sub-confluent monolayers of DBT cells were treated with 0 or 100 U/ml mouse IFN-β for 18 h prior to being infected with virus at an MOI of 1 PFU/cell at 37°C for 45 min. After incubation, inocula were removed, cells were washed with PBS, and fresh medium was added. At 12 h post-infection, cell culture supernatants were removed and 100 μl of supernatant was added to 900 μl TRIzol reagent. Viral genome RNA was purified and the number of viral genome RNA copies present relative to an RNA standard curve were determined as described above. Based on the number of viral genome RNA copies determined by qRT-PCR, an equal number of virus particles from each virus and each condition were combined with an equal number of WT silent or ExoN(-) silent virus particles, respectively. WT silent and ExoN(-) silent viruses were engineered to harbor 10 silent mutations in the probe-binding region of nsp2, allowing separate detection from WT-MHV or ExoN(-) virus genomes, respectively, using a separate probe upon co-infection (24). Next, a fresh, sub-confluent monolayer of DBT cells were co-infected with each combination of viruses at 37°C for 45 min. After incubation, inocula were removed, cells were washed with PBS, and fresh medium was added. At 24 h post-infection, cell culture supernatants were removed and 100 μl of supernatant was added to 900 μl TRIzol reagent. Viral genome RNA was purified. The number of viral genome RNA copies of both reference and silent viruses were determined relative to the appropriate standard curve. The number of viral genome RNA copies relative to the number of silent virus genome RNA copies was determined for each virus and condition. Values are reported as the change in fitness relative to the silent virus.

### Statistical analysis

Statistical tests were applied as noted in the respective figure legends and were determined using GraphPad Prism 6 software (La Jolla, CA).

## ACKNOWLEDGMENTS

We thank members of the Vanderbilt Technologies for Advanced Genomics (VANTAGE) core for rRNA integrity determination services. We thank fellow members of the Denison and Weiss laboratories, specifically Maria Agostini, for helpful discussions. This work was supported by Public Health Service awards T32 HL07751 (J.B.C.) from the National Heart, Lung, and Blood Institute, R01 AI108197 (M.R.D.), and R01 AI104887 (S.R.W.) from the National Institute of Allergy and Infectious Diseases. Additional support was provided by the Elizabeth B. Lamb Center for Pediatric Research.

